# Molecular Property Diagnostic Suite for COVID-19 (MPDS^COVID-19^): An open access disease specific drug discovery portal

**DOI:** 10.1101/2023.08.29.555437

**Authors:** Lipsa Priyadarsinee, Esther Jamir, Selvaraman Nagamani, Hridoy Jyoti Mahanta, Nandan Kumar, Lijo John, Himakshi Sarma, Asheesh Kumar, Anamika Singh Gaur, Rosaleen Sahoo, S. Vaikundamani, N. Arul Murugan, U. Deva Priyakumar, G.P.S. Raghava, Prasad V. Bharatam, Ramakrishnan Parthasarathi, V. Subramanian, G. Madhavi Sastry, G. Narahari Sastry

## Abstract

Computational drug discovery is intrinsically interdisciplinary and has to deal with the multifarious factors which are often dependent on the type of disease. Molecular Property Diagnostic Suite (MPDS) is a Galaxy based web portal which was conceived and developed as a disease specific web portal, originally developed for tuberculosis (MPDS^TB^). As specific computational tools are often required for a given disease, developing a disease specific web portal is highly desirable. This paper emphasises on the development of the customised web portal for COVID-19 infection and is referred to as MPDS^COVID-19^. Expectedly, the MPDS suites of programs have modules which are essentially independent of a given disease, whereas some modules are specific to a particular disease. In the MPDS^COVID-19^ portal, there are modules which are specific to COVID-19, and these are clubbed in SARS-COV-2 disease library. Further, the new additions and/or significant improvements were made to the disease independent modules, besides the addition of tools from galaxy toolshed. This manuscript provides a latest update on the disease independent modules of MPDS after almost 6 years, as well as provide the contemporary information and tool-shed necessary to engage in the drug discovery research of COVID-19. The disease independent modules include file format converter and descriptor calculation under the data processing module; QSAR, pharmacophore, scaffold analysis, active site analysis, docking, screening, drug repurposing tool, virtual screening, visualisation, sequence alignment, phylogenetic analysis under the data analysis module; and various machine learning packages, algorithms and in-house developed machine learning antiviral prediction model are available. The MPDS suite of programs are expected to bring a paradigm shift in computational drug discovery, especially in the academic community, guided through a transparent and open innovation approach. The MPDS^COVID-19^ can be accessed at http://mpds.neist.res.in:8085.

## 1. Introduction

The world has witnessed the coronavirus disease-19 (COVID-19) caused by SARS-CoV-2 in the recent past. The SARS-CoV-2 infection resulted as a pandemic and caused severe damage to healthcare and economy worldwide (Li et al., 2021; Markov et al., 2023; Phukon et al., 2022; Mahanta et al., 2022). In the direction of developing new therapeutics and drugs for severe acute respiratory syndrome-coronavirus-2 (SARS-CoV-2), scientists all over the world have made substantial progress in deciphering the pathophysiology of the disease and its prognosis (Holmes et al., 2021). Further important clues at molecular level details on the viral entry, transcription, translation, translocation, and viral interaction with host cell receptors were obtained. This has provided a definitive knowledge on the druggable target protein of the viral genome (Jackson et al., 2022).

The Molecular Property Diagnostic Suite (MPDS) is a Galaxy (Gu et al., 2021; Galaxy community., 2022; Gaur et al., 2023) based open-source computational drug discovery platform. The development of MPDS is a result of our interest in developing open-source disease specific web portals (Gaur et al., 2017 and 2018; Nagamani et al., 2017). Galaxy offers the flexibility for users to deploy tools using multiple programming languages based on their preferences. Computational methods have played strategically significant role to counter the challenges in drug discovery and development parlances. In this manuscript we put forward a disease specific web portal on COVID-19, which is in line with our earlier web portals, namely MPDS^TB^(Tuberculosis) (Gaur et al., 2017) and MPDS^DM^ (Diabetes Mellitus) (Gaur et al., 2018). Galaxy is an open source, transparent, platform for the bioinformatics, cheminformatics, genomics, proteomics and CADD analysis. Drug discovery involves finding hits converted them into lead and optimising the leads (Badrinarayan et al., 2011). In general, for the target identification and validation, virtual screening (Choudhury et al., 2019), drug design and optimization, ADME-Tox prediction, drug-drug interactions and personalized medicines different computational methods have been employed that leads to more informed decision-making during drug discovery. The intervention of computational approaches was emphatic during pandemics or large-scale infections, as evidenced by the HIV drug discovery in 90’s and COVID-19 drug and vaccine discovery during the recent pandemic.

Development of customized computational tools and software is essential to comply with the recent advances in computer science and data analytics. Considering the enormous differences among diseases, it has been felt that each disease may have a specific requirement from the (computational) drug design and discovery viewpoint (Jackso et al., 2022; Gu et al., 2021; Galaxy Community., 2022). MPDS^COVID-19^ is a customized web portal (Fig. 1) that is designed to address the questions pertaining to computational drug discovery for COVID-19, and accordingly, a large number of modules have been integrated. Among these modules, some of them are specific to COVID-19, while others are essentially disease independent. Galaxy is an open-source web-based platform that provides access to various tools, huge amounts of data that can be integrated from different fields such as bioinformatics, cheminformatics, genomics, transcriptomics, and used to develop models based on advanced artificial intelligence (AI) and machine learning (ML) approaches (Gu et al., 2021; Galaxy Community., 2022; Gaur et al., 2023). The advantage of this platform is that the source code is publicly available and the users can customize, scale-up the platform to their interest by integrating multiple tools and also design their own workflows. The transparency, reproducibility, and accessibility of this server made the developers easy to integrate their scripts, tools and codes in an efficient manner. A list of critical features of the MPDS^COVID-19^ portal along with a brief justification is provided below.

**Fig. 1.**
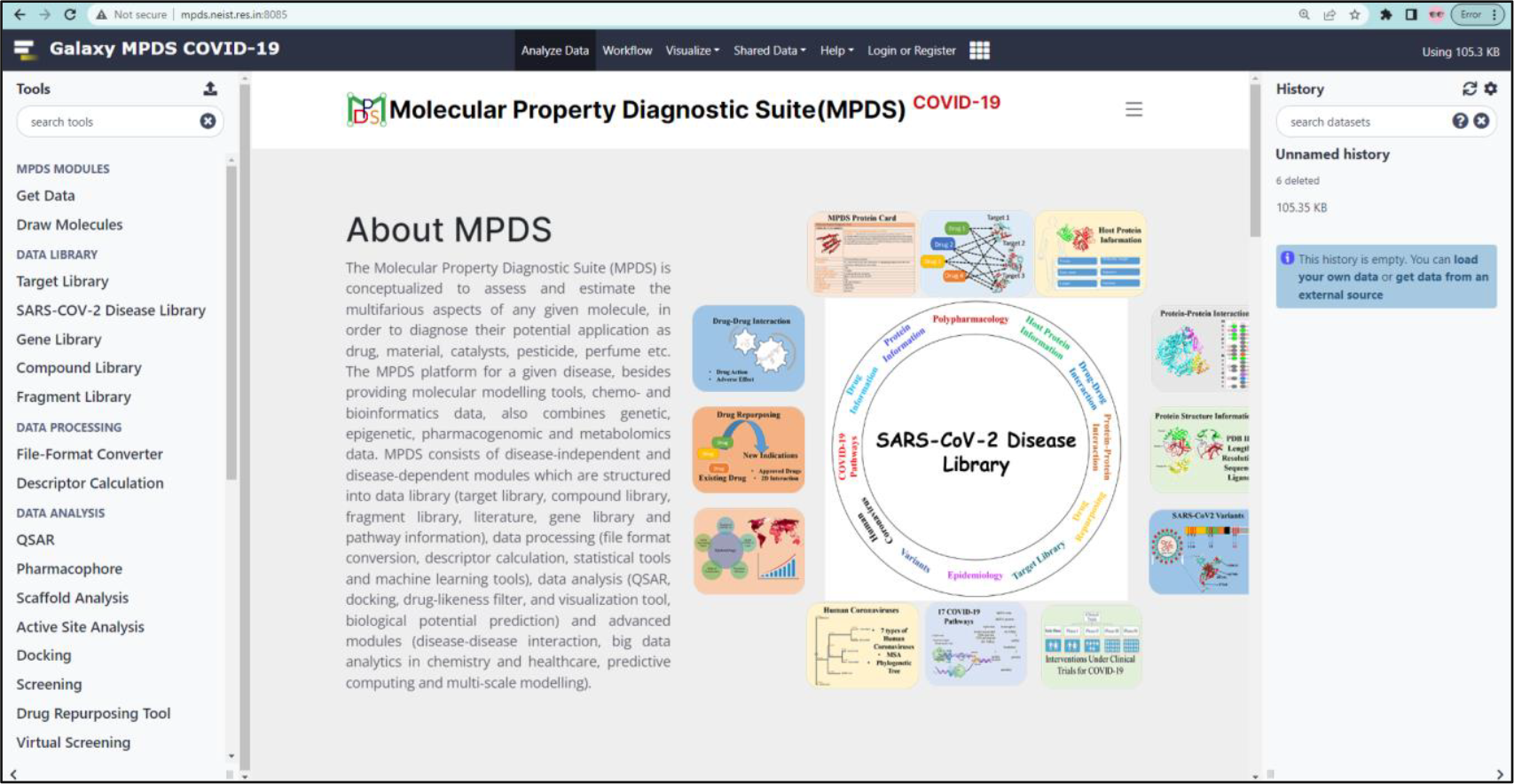
The home page of MPDS^COVID19^. The portal can be accessed at http://mpds.neist.res.in:8085. The MPDS^COVID19^ has structured as *Data library, Data processing, Data analysis and Advanced Modules*.

### Integration

his portal effectively integrates the molecular modelling, informatics, simulations, literature survey and prior art on the disease pathophysiology and drug discovery.

### Latest literature and updates

One of the important aspects of developing the disease specific web portals is to provide the latest information on the development of drugs, related literature and all the information pertaining to the disease. It is also to be noted that any particular *in silico* script, algorithm, database, web portal or software which may be of use can easily be integrated with the MPDS^COVID-19^.

### Easy access of data

The web portal contains both disease dependent as well as disease independent modules which makes it easy to access the desired data of the particular disease and perform the desired analysis using the available tools.

### Data collection

The information is collected from known databases/servers such as Protein Data Bank (PDB) (Sussman et al., 1998), PubMed (Canese et al., 2013), UniProt (Uniprot Consortium., 2019), DrugBank (Wishart et al., 2008), DrugCentral (Ursu et al., 2017), World Health Organization (WHO) (https://www.who.int/emergencies/diseases/novel-coronavirus-2019) and other open access databases (Supplementary Table S1) which make it easy for the users to follow-up.

### Interface

This suite has an advantage of user-friendly interface; Galaxy, allows the users to modify, add any desired tools or data with the permission of the developer.

### Uniqueness

This platform allows one to do computational drug discovery without using any computing resources from the user side. Internet connection appear to be the only requirement and then, the whole computing loads are taken care by the MPDS local server. MPDS^COVID-19^ serves as an efficient platform for conducting virtual screening (for COVID-19 associated targets), pharmacophore modelling, and developing ML models and 2D-QSAR.

Approaches employed in the area of drug discovery, and especially the *in-silico* methods are going to witness tremendous changes owing to the unprecedented progress in the hardware, software as well as AI and internet of things (IoT) based methods (Supplementary Table S2) (Murugan et al., 2022). Trying to break the existing paradigms in the computational drug discovery, by developing a disease specific holistic portal, integrating the existing knowledge with the computational modelling appears to be promising. To this end, our attempts of developing a series of disease specific MPDS web portals may drive the research community to share the program and software for freely.

## 2. Materials and methods

### 2.1. The Galaxy architecture

Galaxy (https://galaxyproject.org/) (Galaxy Community., 2022) is a web-based application that can be used to create user-defined workflows and is readily intended to interact with any other software or script. The platform has been effectively used by a number of scientists worldwide to analyse the biomedical data available in the various field of genomics, proteomics, transcriptomics, bioinformatics and many more (Supplementary Fig. S2). Galaxy platform continues to focus on various aspects viz. available data analysis along with the tool developer to integrate their tools, reproducible analysis by using the specific platform and making the platform simple by transparent communication to reuse and extend the availability. The galaxy platform consists of four main components such as a) the main public galaxy server (https://usegalaxy.org); b) Galaxy framework and software ecosystem (https://github.com/galaxyproject); c) the galaxy toolshed; and d) the galaxy community. The list of available servers and web portals deployed using Galaxy platform are listed in (Supplementary Table S3), along with their URL and use. The detailed information and architecture of galaxy core components are stated in Fig. 2.

**Fig. 2.**
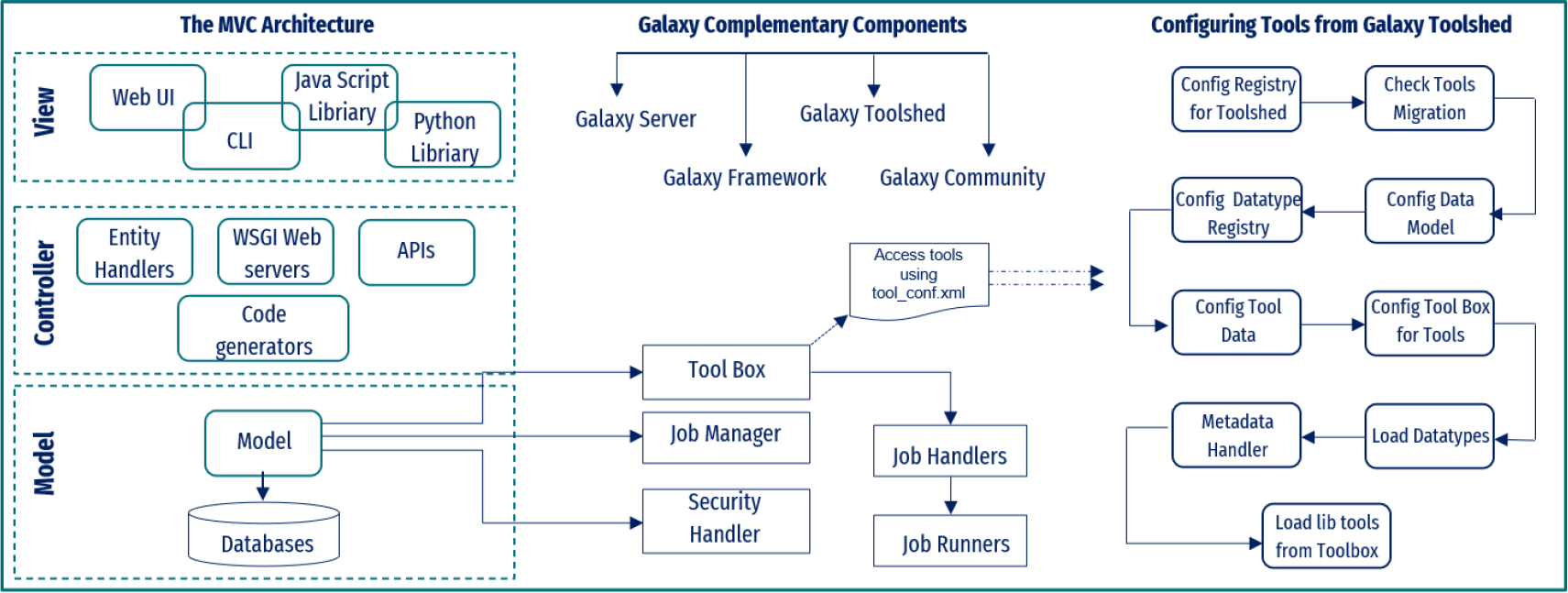
Architecture of Galaxy core components and tool configuration.

### 2.2. Architecture of the MPDS^COVID-19^ software suite

Galaxy application can be installed on UNIX/Linux operating system and it has both frontend and backend with stability and flexibility in tools integration and data analysis. The backend of galaxy is operated by the python server with flexibility and also driven by plugins. The front-end architecture makes the users’ task easy and consistent while the backend works more towards integrating different new technologies (Gaur et al., 2023). In the MPDS^COVID-19^ web portal, the database on COVID-19 has been developed by using in house PHP and MySQL in the backend and integration of new tools and software using in house Perl and Python scripts. The detailed architecture of the MPDS^COVID-19^ is depicted in Fig. 3.

**Fig. 3.**
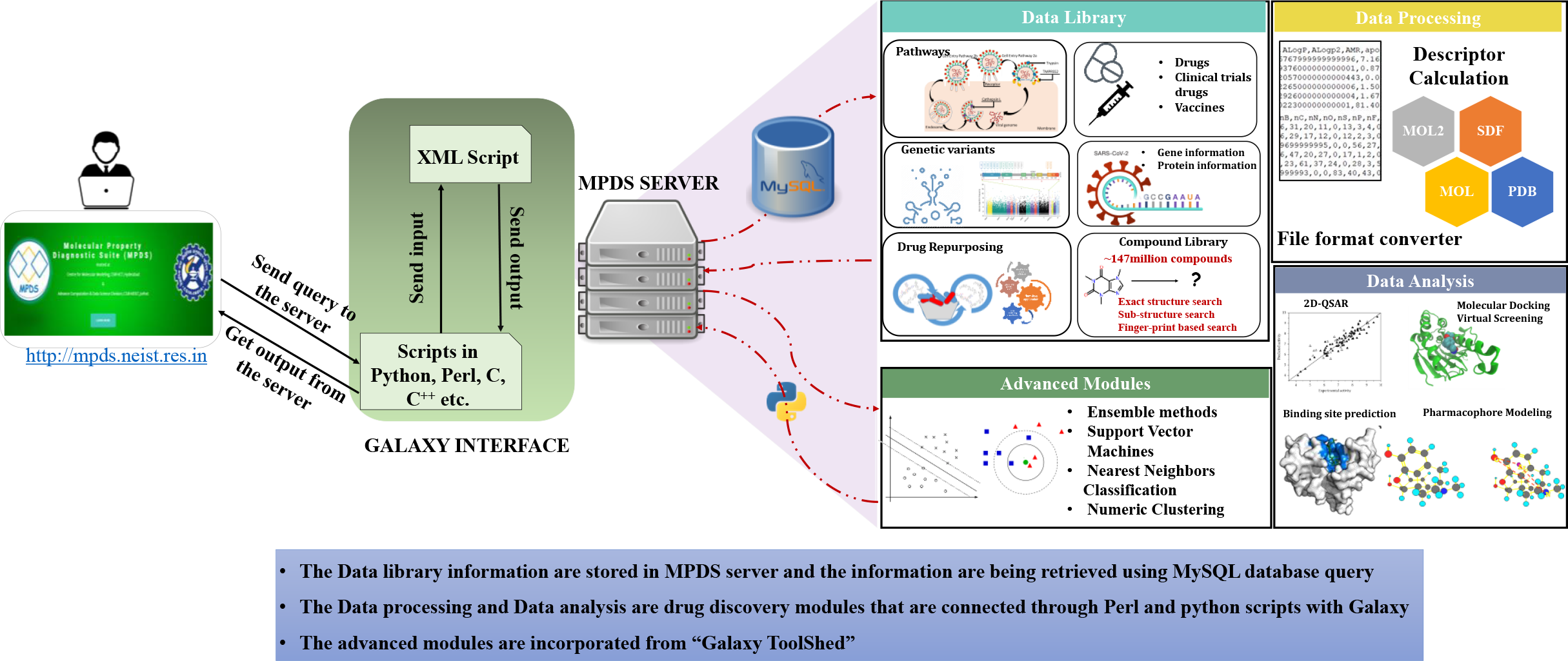
The Galaxy architecture of MPDS^COVID-19^. The users can upload their data (i.e., macro and small molecules) or they can select from MPDS data library. The uploaded or selected information will be available in Galaxy history panel. The users can submit the calculation in the main window and the results will be appeared in the history panel.

The initial design of the MPDS web portal involves three sections: a) Data library, b) Data processing and c) Data analysis. The data library section is generally consisting of disease specific modules and provides the information on SARS-COV-2 Fig.4. Thus, MPDS^COVID-19^ has a large number of SARS-CoV-2 related modules and the web portal also provide links to tools that are specifically designed for antivirals in general and SARS-CoV-2 in particular. The major updates in the data library are the inclusion of fragment library (Gaur et al., 2022) as separate module and a greatly revised form of compound library (John et al., 2023). The data processing module underwent little change expect for the newer versions of file format converter and descriptor calculation. However substantial additions were made in data analysis module Fig. 5. A couple of tools in the predictive module were also developed. A major attractive feature of the MPDS^COVID-19^ is the addition of advanced modules which consists of the employment of machine learning approaches.

### 2.3. Collection of data

The extensive literature survey from the PubMed, google scholar and use of freely accessible databases and tools, colloids various detailed information on the literatures, SARS-CoV-2 targets, gene and their pathway information. Additionally, it also contains the information on repurposed drugs against various targets of SARS-CoV-2, mutational variants, polypharmacology for COVID-19 (Supplementary Fig. S3) (Jamir et al., 2022), drug-drug interaction information, Protein-protein interaction (PPI) study (Sarma et al., 2022a; Sarma et al., 2022b), host protein information, epidemiology and inhibitors database. The extensive information on SARS-CoV-2 proteins were collected from the National Center for Biotechnology Information (NCBI) (Sayer et al., 2018), PDB (Sussman et al., 1998), PubMed (Canese et al., 2013), and Kyoto Encyclopaedia of Genes and Genomes (KEGG) (Kanehisa et al., 2000). The structure information was collected from PDB by using the advanced search query option followed by the create custom report to get the desired information on each protein. The PPI information was collected from various literatures downloaded from PubMed and from online search by using different keywords like “PPI SARS-CoV2 during viral entry”, “PPI During Viral Propagation/survival”, “PPI of Immune evasion” and the other detailed analysis on PPI carried out in our group. The approved drugs, drugs in different phases of clinical trials were collected from the databases such as ChEMBL (Mendez et al., 2019), DrugBank, and PubChem (Kim et al., 2021). Variants, epidemiology information were collected from the WHO (https://www.who.int/activities/tracking-SARS-CoV-2-variants) and European Centre for Disease Prevention and Control (https://www.ecdc.europa.eu/en/covid-19). The gene information contains the sequence, PDB ID, structure, domain information, active site residues, their function along with various available database IDs for the structural, non-structural as well as accessory proteins involved in SARS-CoV-2 (Jamir et al., 2022). The library also provides a detailed description of the pathways involved in COVID-19 infection. It contains information on 9 PPIs that take part in the various pathways starting from viral entry, replication, and transcription to immune invasion (Sarma et al., 2022a; Sarma et al., 2022b). In the drug information, a list of already approved FDA drugs and other emergency and promising drugs for COVID-19 are provided which gives an overview of all the drugs that are possibly available for studying in the drug development process for SARS-CoV-2.

## 3. Results

### 3.1. Development of disease specific web portal

MPDS disease specific portal, developed on the galaxy platform aims to provide the essential information on the existing data in a single platform for easy access to all the users worldwide. The web portal is designed into three modules namely 1. Data Library, 2. Data Processing and Data Analysis.

### 3.2. Disease dependent modules

#### 3.2.1. Module 1: Data library

The Data library comprises of different categories (target library, SARS-CoV-2 disease library and gene library). The target library contains around 3161 PDBs of proteins associated with SARS-CoV-2 with detailed structural information on their molecular weight, structure title, PDB IDs, experimental models, date, resolution, structure title, structure keywords, ligand name and its molecular weight.

#### 3.2.2. Preparation of SARS-CoV-2 disease library

The literature module provides vast information on the pandemic COVID-19 outbreak, druggable targets, the epidemiology study in various regions, and geographical mapping of coronavirus disease worldwide. As a part of these disease specific modules, the portal contains the libraries as discussed in section 2.3. The protein information contains sequence and structural information related to 4 structural, 16 non-structural and 9 accessory proteins of SARS-CoV-2. Importantly, users can access protein information such as the sequence, 3D structure of protein, their functions, amino acid length, molecular weight, protein domain information, active site residues, selected PDB ID, NCBI ID, UNIPROT ID, PUBMED ID for literature and KEGG pathway ID. Information on different mutational variants of SARS-CoV-2 such as variants of interest (VOIs), variants of concern (VOCs), previously circulating (VOIs), previously circulating (VOCs) and formerly mentioned variants (FMVs) for spike protein, and the epidemiology information are incorporated in the disease library.

The MPDS^COVID-19^ has also 3D structures of SARS-CoV-2 inhibitors database for different targets. Other than this, the clinical trial dataset, cell experiment dataset and virtual screening dataset are available in this section. The dataset was collected and curated from the published literature. The database contains inhibitors along with the binding affinity from ChEMBL [29] and Binding database (Liu et al., 2007). The users can download the 3D structures of the 7721 molecules for different computational drug discovery calculations. The visual representation of the SARS-CoV-2 disease library is illustrated in Fig. 4.

**Fig. 4.**
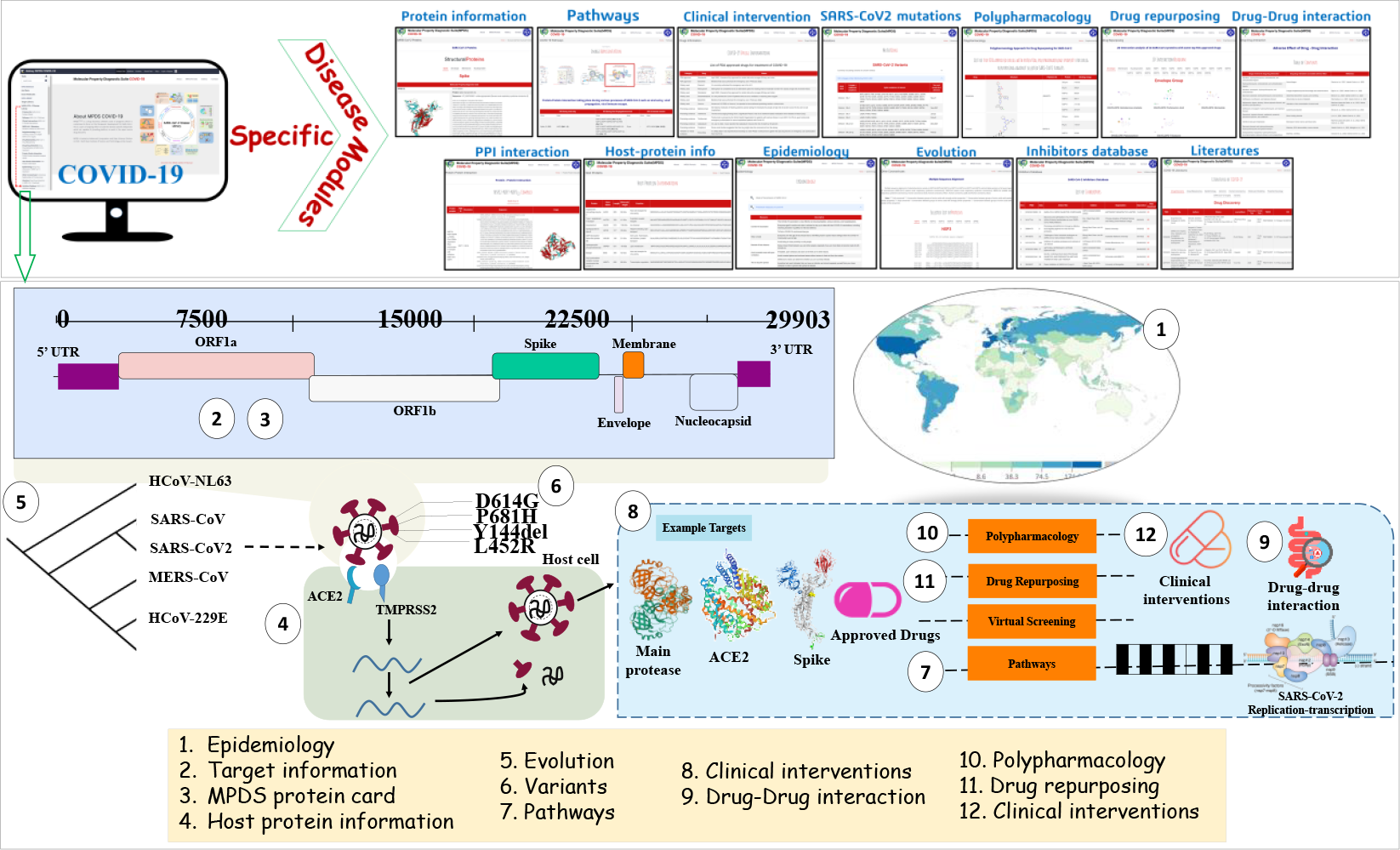
The diagrammatic representation depicting incorporated various disease specific information in MPDS^COVID-19^ disease library.

#### 3.2.3. Gene library

The gene library in MPDS^COVID-19^ contains a unique identifier called as gene ID. Upon searching for the gene ID, the corresponding gene card will be generated, and it contain, gene name, gene description, the gene ID and characterization.

### 3.3. Disease independent modules

In addition to the data library module in the disease dependent category, there are several disease independent modules, as described in the following sections.

#### 3.3.1. Compound and Fragment library

The compound library module in the initial MPDS suite, has about 110 million compounds, where the chemical space was divided into classes. Each class represent a unique structural feature and to limit the number of compounds in a group, the number of molecules in a class was divided into clusters. Recent version of MPDS compound library has over 149 million unique molecules and distributed into 57 classes. Every molecule is provided with an MPDS-AadharID, which unambiguously characterize the molecule by assigning a unique class and cluster number (John et al., 2023). Users can submit the data in a variety of forms, including SMILES, InChIKey, mol, mol2 and sdf formats. If the query molecule exists in the database, an MPDS-AadharID is generated. In case the query molecule is not available in the database, a message will be displayed indicating that the molecule does not exist in the database. In such a case the user may contact the manager of webserver, and a new MPDS-AadharID number will be generated. Recently, fragment library module of the MPDS has been added (Gaur et al., 2022). The fragment library, contains 107,614 unique fragments which may be further classified as rings, linkers, and substituents.

#### 3.3.2. Data processing

Different tools have unique requirements for the input files and produce the output files in another file format. In order to fulfil the requirements of different tools, various file format converter tools have been used from the galaxy toolshed for inter conversion among various file formats. There are many input/output formats, viz., .sdf, .mol, .mol2, .cml, InChI, SMILES, and .pdb; and correspondingly the output formats (.mol2, .sdf, .mol and .pdb) are incorporated into the MPDS. The module also includes a tool for converting the 2D structure to 3D structure of compounds. MPDS provides two descriptor calculation tools namely PaDEL and CDK to calculate different 2D and 3D properties of the compounds.

#### 3.3.3. Data analysis

The data analysis modules include data mining, and quantitative structure activity relationship (QSAR) (Srivastava et al., 2012) through various galaxy implemented tools such as McQSAR, wekatool and SVMlight. Expectedly, a large number of computational tools have been developed in the area of drug discovery, and covering all of them is a daunting task and outside the purview of this portal. However, we made an attempt to integrate the available open-source software packages and link them in our web portal. Besides, here we provide a brief description of the selected data analysis tools that may be of high practical utility for the people desirous of using our web portal. The modules include a) Pharmacophore (Choudhury et al., 2016), b) Scaffold analysis, c) Active site analysis, d) Docking, e) Screening, f) Drug repurposing tool (Lagunin et al., 2000), g) Virtual screening, h) Visualization, i) Sequence alignment, and the advanced module contains various machine learning tools along with the inhouse developed machine learning antiviral prediction model (John et al., 2022). The disease independent modules are depicted in Fig. 5.

**Fig. 5.**
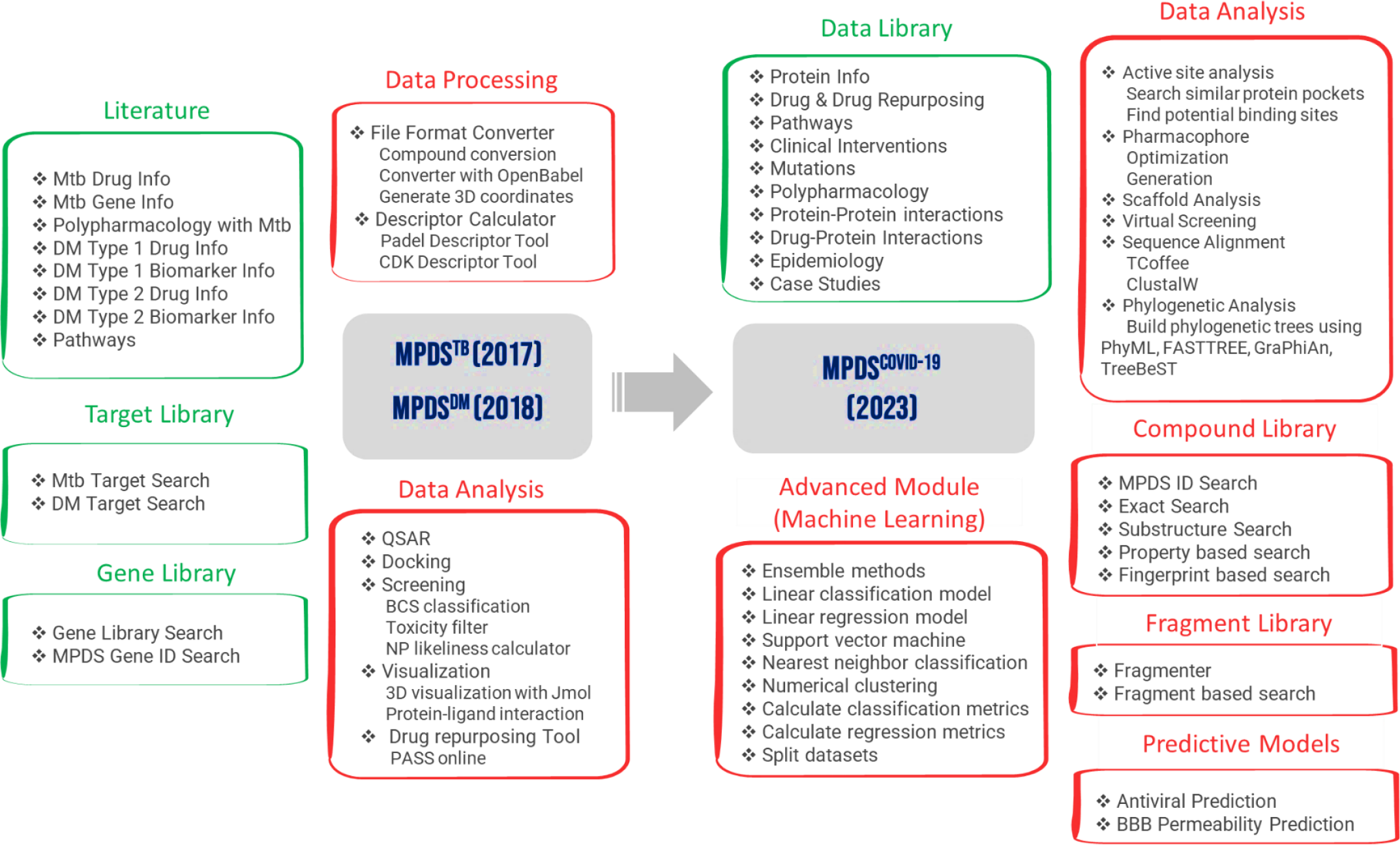
Evolution of MPDS^COVID-19^ portal, in terms of time and features, with integration of new disease dependent modules (shown in green headers and boxes) and independent modules (shown in red headers and boxes). The MPDS^TB^ and MPDS^DM^ were developed in Galaxy 16.01 (released in 2016) version while MPDSCovid-19 has been developed with Galaxy 19.05 (released in 2019). There were 35 tools from Galaxy Toolshed which were integrated in MPDS^TB^ and MPDS^DM^ whereas in MPDS^COVID-19^, 59 tools from Galaxy Toolshed have been integrated. Compound Library (http://mpds.neist.res.in:8086/), Antiviral Prediction Tool (http://acds.neist.res.in:8501/) and Fragment Library are tools/libraries developed by the group and have been integrated into MPDS^COVID-19^. All modules shown on MPDS^COVID-19^ are the additions that were done to the earlier versions in 2017 and 2018.

## 4. Discussion

MPDS^COVID-19^ software suite has a lot of important data that are collected along with the molecular modelling and software tools that are implemented. Besides, the web portal effectively integrates several modules that are developed in the areas of cheminformatics and CADD which are available in the Galaxy portal. All the data are extracted from the research papers and the freely available online platforms (Supplementary Table S1) and manually curated before incorporating in the database. This effort is for the future preparedness of the drug discovery software, which essentially focuses on the larger problem and needs to be addressed. This is in contrast with the conventional approaches of developing the software, where the focus is on developing the methods and their implementation in an effective way.

This change in the paradigm warrants a very effective integration of the AI/ML and IoT based techniques to the molecular modelling, bioinformatics and cheminformatics approaches. The conventional molecular modelling techniques are based on the principle of classical mechanics and quantum mechanics or statistical mechanics. In contrast the data driven approaches are entirely heuristic and thereby adopts a highly complementary approach compare to molecular modelling. Therefore, an effective integration of these two approaches is indispensable for obtaining deeper insights into the problems of drug design.

### 4.1. Tips to use MPDS^COVID-19^

The MPDS galaxy user server has been installed at CSIR-NEIST, Jorhat and it can be accessed at http://mpds.neist.res.in:8085/. However, useful data can be obtained such as drug discovery calculations, retrieval of information, QSAR models and the generation of workflows. One may refer to the manual which is available in the website, for the usage of several modules and to access the SARS-CoV-2 disease library. The software suite enables users to submit data from their local computer and freely access and download the outcomes. With the authorization of the administrator, users are permitted to alter and include the necessary tools, databases, and software of interest. Logging into the MPDS server at the host institute might provide a lot of useful information and depends on the traffic and network speed and connectivity issues Fig. 6.

**Fig. 6.**
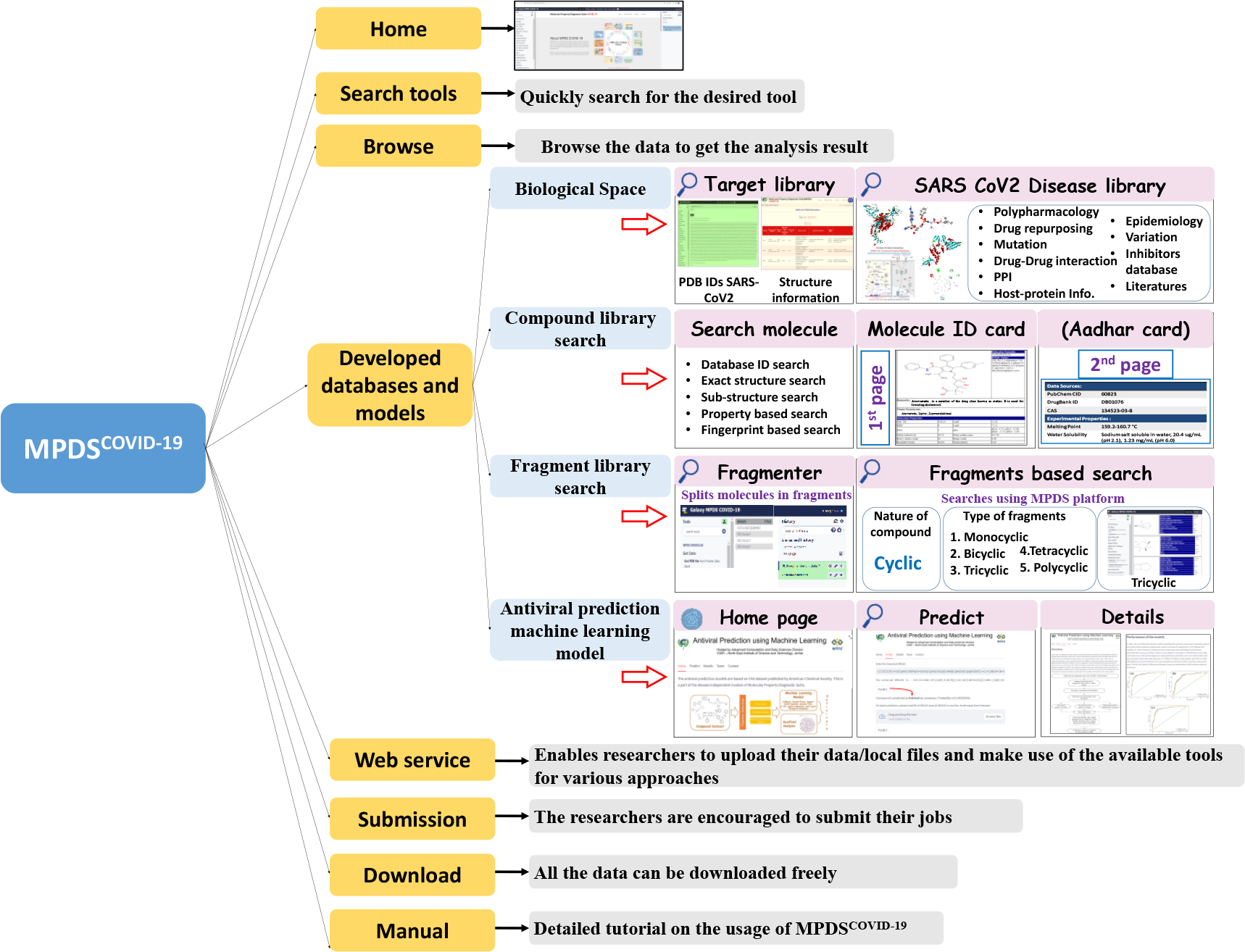
The scheme of the architecture of data integration and manual curation along with the use of developed compound library, fragment library and antiviral prediction machine learning model.

A straightforward approach to evaluate the utility of any tool or software is to try some case studies. We provide about 6 case studies and describe them in detail in the supporting information as well as in the manual (Supplementary Figs. S1a to 1f).

### 4.2. Comparison with other resources

There are several resources in the field of drug discovery, such as OSDD platform, ODDT (Wojcikowski et al., 2015), SCFBio software, CYCLICA, Orion-cloud-based platform, Gromacs (Abraham et al., 2015), Amber (Case et al., 2005), AutoDock (Morris et al., 1991), AutoDock Vina (Trott et al., 2010), Schrodinger Suite (Bhachoo et al., 2017), which are widely used for drug discovery solutions. Several databases and tools have been developed specifically for COVID-19 drug discovery. The databases include covid-evidence, OxCOVID19, LitCovid, DockCoV2, SARS-CoV-2, COVID-19 data portal, COVIDium, SCoV2-MD and additional databases. The detailed information on the available databases on drug discovery for COVID-19 are listed in (Supplementary Table S4). Comparatively, MPDS is a comprehensive suite of databases, tools and software developed on the open-source Galaxy platform that combines all elements of the drug discovery process into one user-friendly platform. Portals specific to different diseases are included in the suite. The MPDS^COVID-19^ contains all manually curated disease specific information on SARS-CoV-2.

Upon realization of the importance of ADMET, solubility, selectivity, and consideration of risk to benefit ratio due to the side effects, more profound and challenging questions have emerged. The tremendous growth in the generation of data resulted in bigger challenges as well as the opportunities to push the drug discovery efforts to all together to a different plane. Thus, looking forward to getting answer by the integration of conventional molecular modelling approaches with that of AI, ML and IoT based methods are essential.

While talking about the advancement in AI and ML, it also comes with certain limitations. The AI/ML algorithms mainly rely on the large set of accurately curated data sets for further training and validation. In case of drug discovery, it is challenging to get well curated data when it comes about the rare diseases or novel targets (Dong et al., 2022). Furthermore, there may be the chances of overfitting in case of machine learning model, limitations in the chemical space exploration since the novel drug candidates or unconventional chemical structures may not be well captured by the trained AI/ML models (Gysi et al., 2023).

## 5. Conclusions

The objective of this work is to develop an open-source computational drug discovery platform for COVID-19. *MPDS disease specific web portal is an initiative which is thoughtfully customized one-stop solution*, has a great potential to drive the open-source computational drug discovery. Diseases are different, in terms of their pathophysiology and manifestation and therefore developing a customized approach is the need of the hour. Especially, in case of COVID-19, a great deal of data has been generated in a quick time owing to the unprecedented impact of the pandemic necessitating capture of all that data. The disease independent modules can be of great value for the researchers, especially academicians without access to the commercial software, engaged in the drug discovery research. In order to make the research more transparent and effective, the open-source platforms should emerge as the formidable alternatives to the commercial computational drug discovery packages (Murtazalieva et al., 2017; Druzhilovskiy et al., 2017).

The disease dependent modules will provide a right platform to ask profound questions related to a disease, and some of them can be quite focused or even provocative: a) Is it possible to synthesize personalized medicines? b) Can we understand the altered pathophysiology of a given disease in light of comorbidity, which warrants to avoid/employ specific drugs? c) Can we understand drug-drug interactions and come up with recommendations for administering them in cases where a patient has more than one disease? MPDS is one of the approaches which has the potential to formulate and address such questions, by augmenting new algorithms and modules.

## CRediT authorship contribution statement

The architecture and content of the software are conceived by GNS. All authors have contributed to incorporate the data in data library and development of the portal. The manuscript has been proof-read by all the authors.

## Acknowledgements

DBT is thanked for the financial support in the form of Centre of Excellence in Advanced Computation and Data Sciences (Ref. No: BT/PR40188/BTIS/137/27/2021). CSIR is thanked for supporting the research in the form of project HCP41.

