## Supplementary Material for "Molecular Property Diagnostic Suite for COVID-19 (MPDS^COVID-19^): An open access disease specific drug discovery portal"

**Figure S1a: Case study 1-** Screening of phytochemicals for antiviral activity against SARS-CoV2 targets.

**Figure S1b: Case study 2-** Drug repurposing for SARS-CoV2 protein NSP5 through virtual screening

**Figure S1c: Case study 3-**Screening of phytochemicals for natural product likeness properties and docking against SARS-CoV2 protein involved in viral entry

**Figure S1d: Case study 4-** Phylogenetic tree generation of seven types of Human Coronavirus using the available sequence from the SARS-CoV2 disease library

**Figure S1e: Case study 5-** Perform QSAR calculation of FDA approved drugs using the available drugs from the SARS-CoV2 disease library

**Figure S1f: Case study 6-** Investigating Repurposed Drugs' Binding Interactions with SARS-CoV-2 RdRp Using MPDS-COVID Docking

**Figure S2.** An illustration of various tools available in the Galaxy platform under different categories focused in the field of drug discovery.

**Figure S3.** An overview of drug repurposing approach for COVID-19 disease which involves identifying of existing drugs with potential therapeutic indications against multiple targets. This may include assessing potential novel uses for drugs based on their known mechanisms of action and interactions with druggable targets, as well as screening existing drugs against the targets in the same pathway or targets from multiple pathways. Applying a unique combination of drug repurposing and polypharmacology approach has led to identification of potential leads that can be repurposed for COVID-19 disease.

**Table S1.** List of AI-Powered drug discovery tools along with its URL, brief description and reference.

**Table S2.** List of available severs and web portals deployed using Galaxy platform.

**Table S3.** List of publicly available various COVID-19 disease related databases and their description.

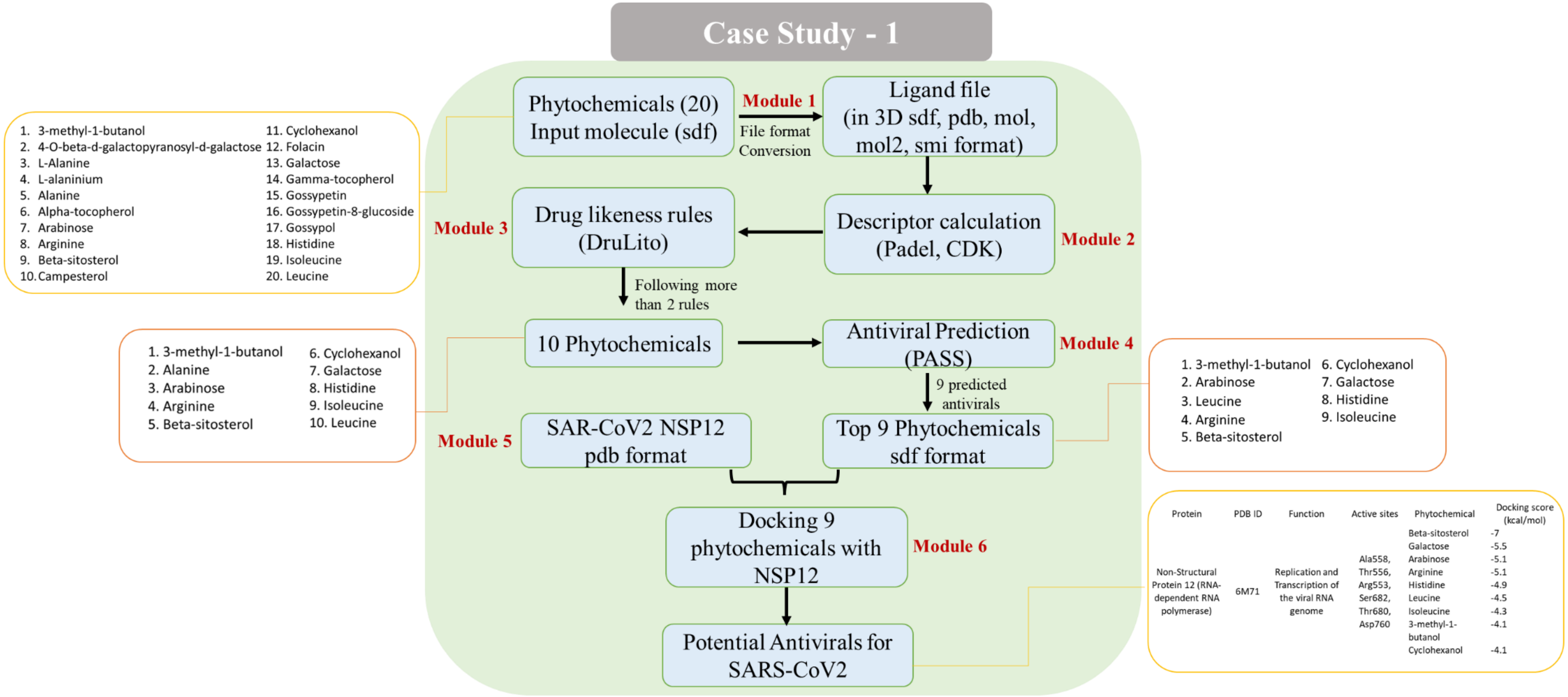


**Figure S1a**: Screening of phytochemicals for antiviral activity against SARS-CoV2 targets

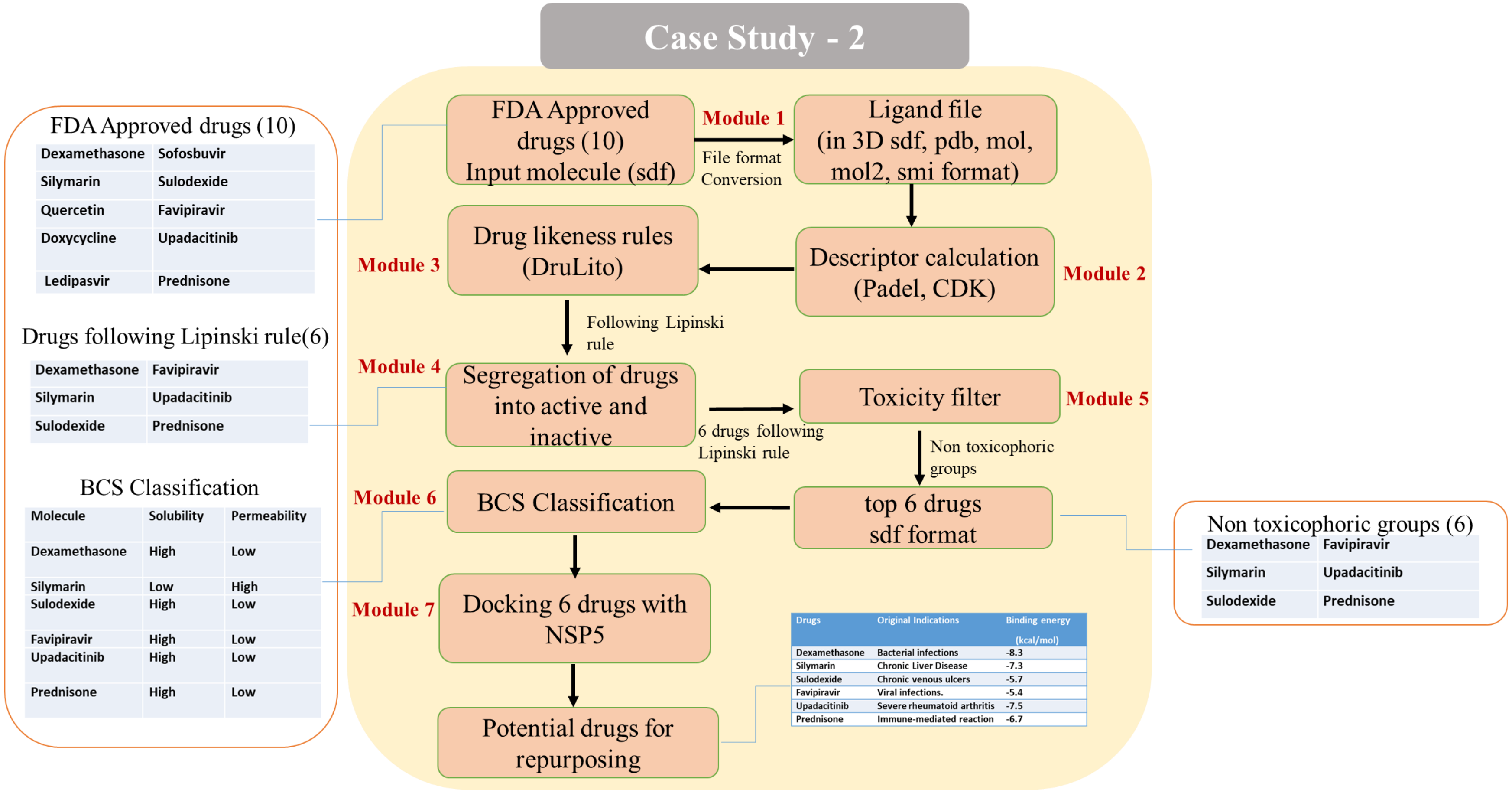


**Figure S1b:** Drug repurposing for SARS-CoV2 protein NSP5 through virtual screening

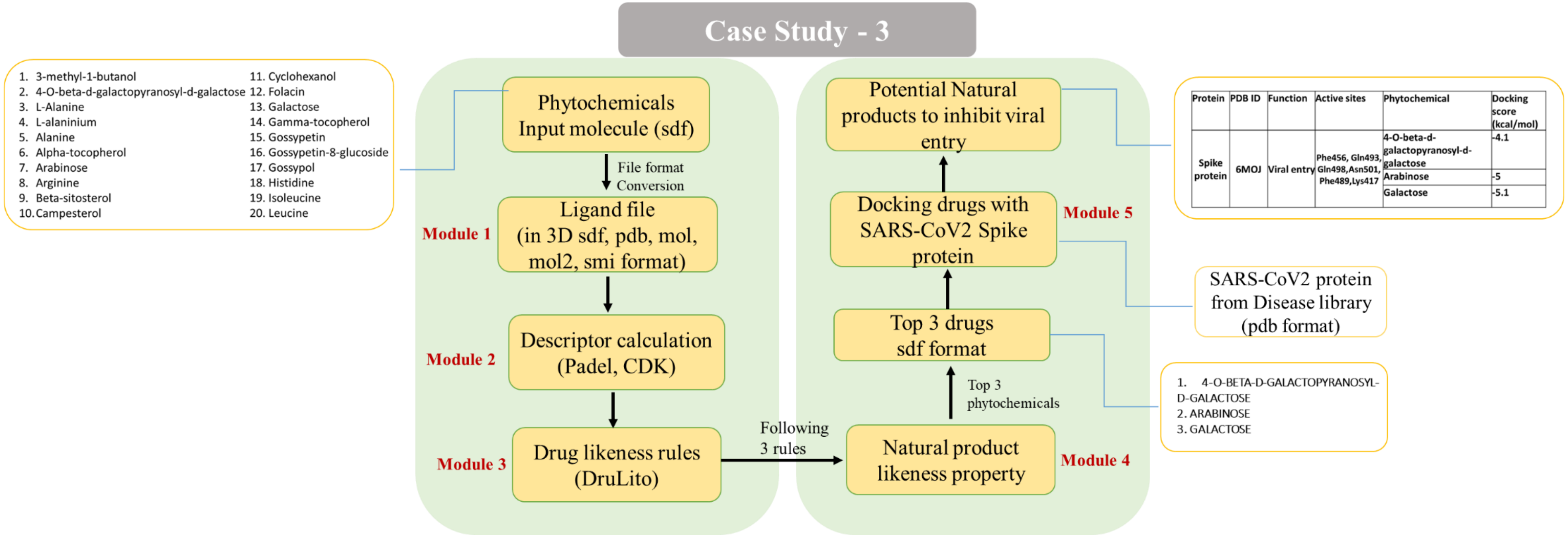


**Figure S1c:** Screening of phytochemicals for natural product likeness properties and docking against SARS-CoV2 protein involved in viral entry

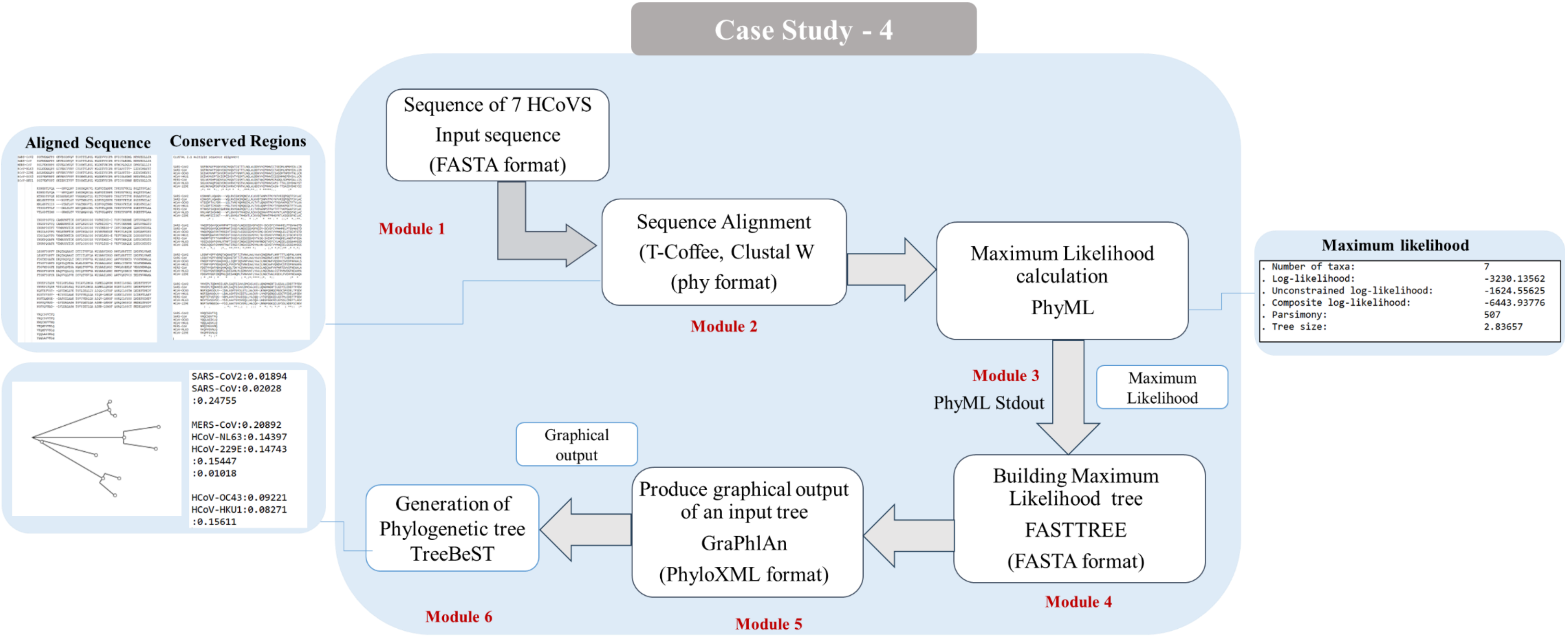


**Figure S1d:** Phylogenetic tree generation of seven types of Human Coronavirus using the available sequence from the SARS-CoV2 disease library

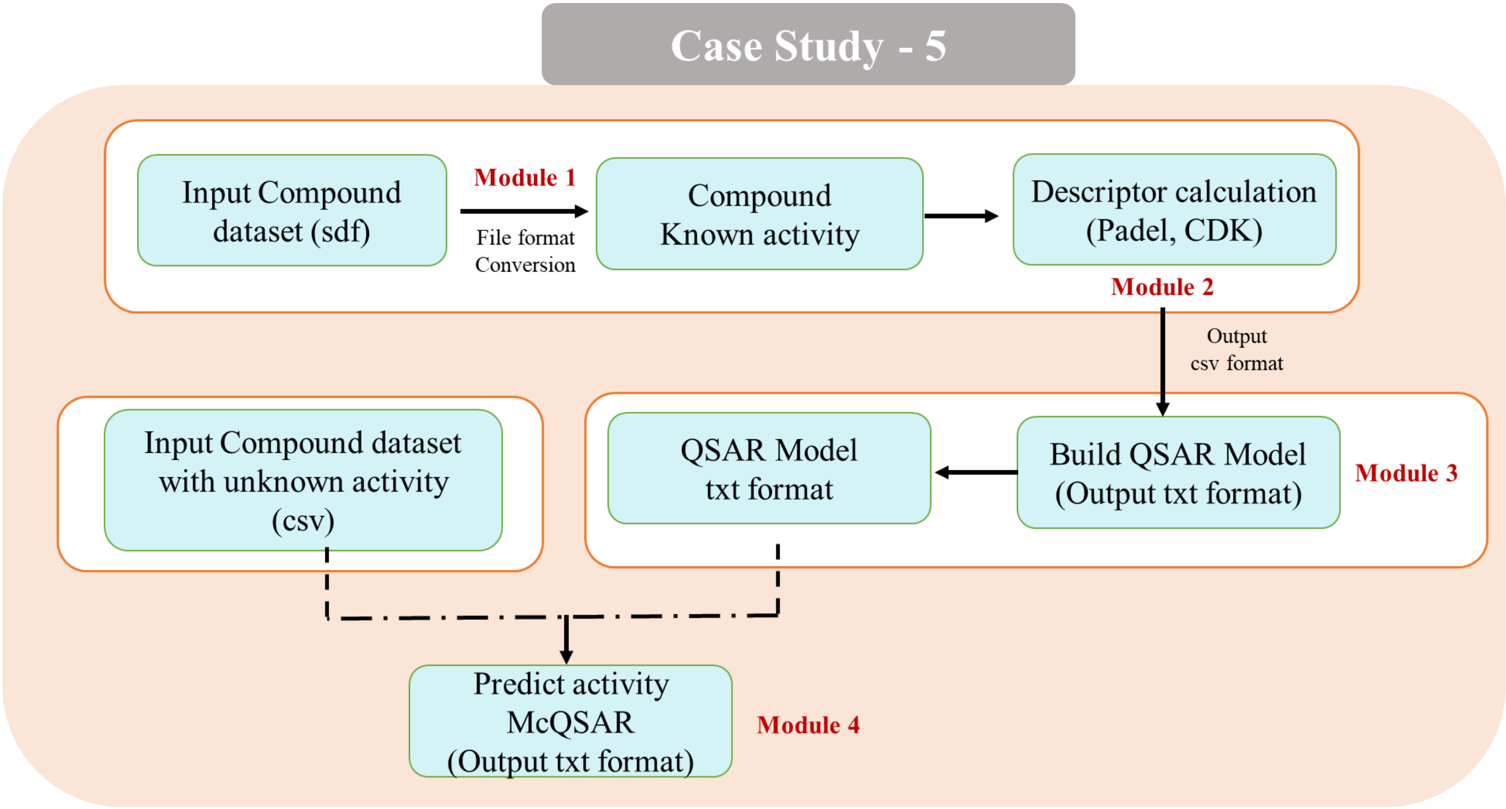


**Figure S1e:** Perform QSAR calculation of FDA approved drugs using the available drugs from the SARS-CoV2 disease library

### Case Study-6

**Investigating Repurposed Drugs' Binding Interactions with SARS-CoV-2 RdRp Using MPDS-COVID Docking**

This study is to demonstrate the potential of the MPDS-COVID docking module in facilitating the development of multi-targeted therapies against SARS-CoV-2. This study showcases the efficacy of MPDS-COVID docking module in examining how repurposed drugs interact with SARS-CoV-2's RNA-dependent RNA polymerase (RdRp).

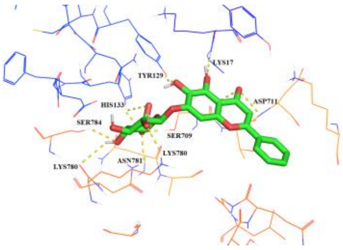


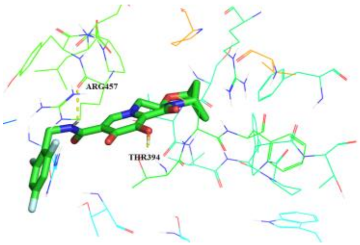


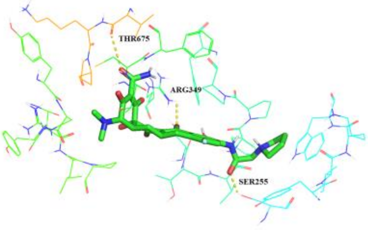


**Baicalin** **Bictegravir** **Eravacycline**

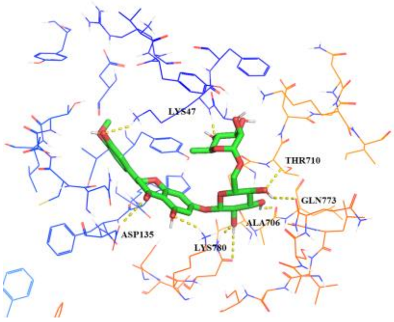


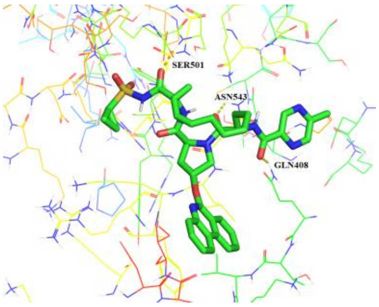


**Hesperidin** **Paritaprevir**

| Sl. No. | Drug Name | Binding Affinity (kcal/mol) | Interacting Residues |
|---|---|---|---|
| 1. | Baicalin | -8.7 | LYS17, TYR129, HIS133, SER709, ASP711, LYS780, **LYS780**, ASN781, SER784 |
| 2. | Hesperidin | -9.7 | LYS47, ASP135, ALA706, THR710, GLN773, **LYS780** |
| 3. | Bictegravir | -8.7 | THR394, ARG457 |
| 4. | Eravacycline | -8.5 | SER255, ARG349, THR675 |
| 5. | Paritaprevir | -10.1 | GLN408, SER501, ASN543 |

**Figure S1f:** Binding affinities and interacting residues of repurposed drug molecules with the SARS-CoV-2 RNA-dependent RNA polymerase (PDB ID: 6M71). The MPDS-COVID docking module utilized specific parameters for the docking process.

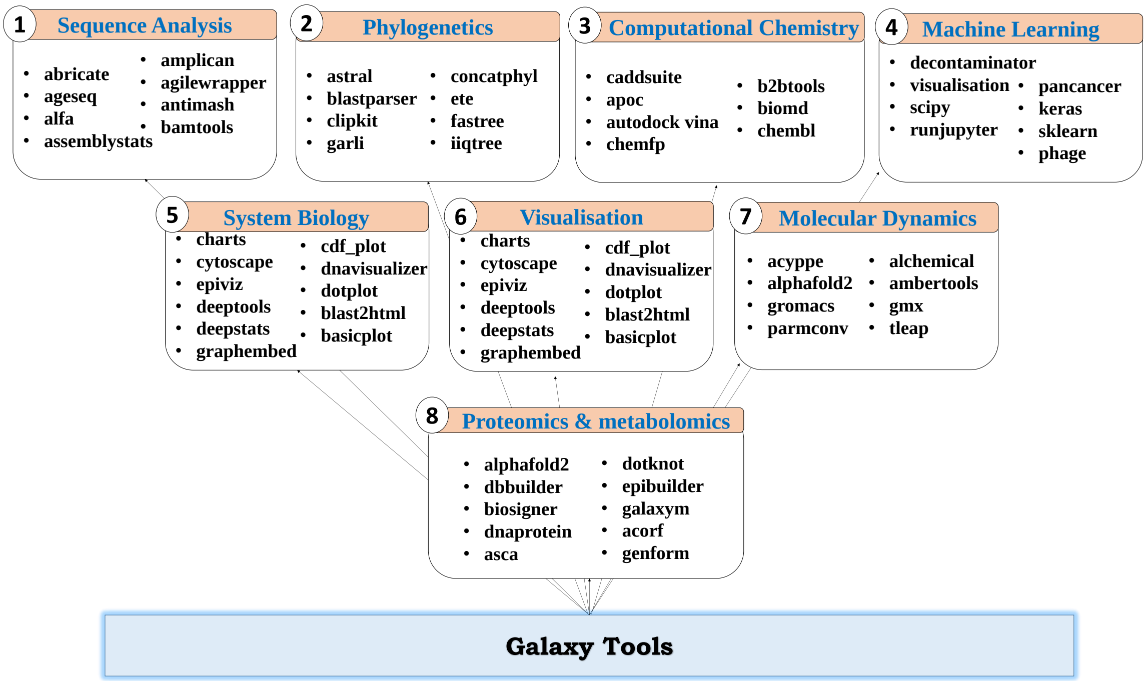


**Figure S2.** An illustration of various tools available in the Galaxy platform under different categories focused in the field of drug discovery.

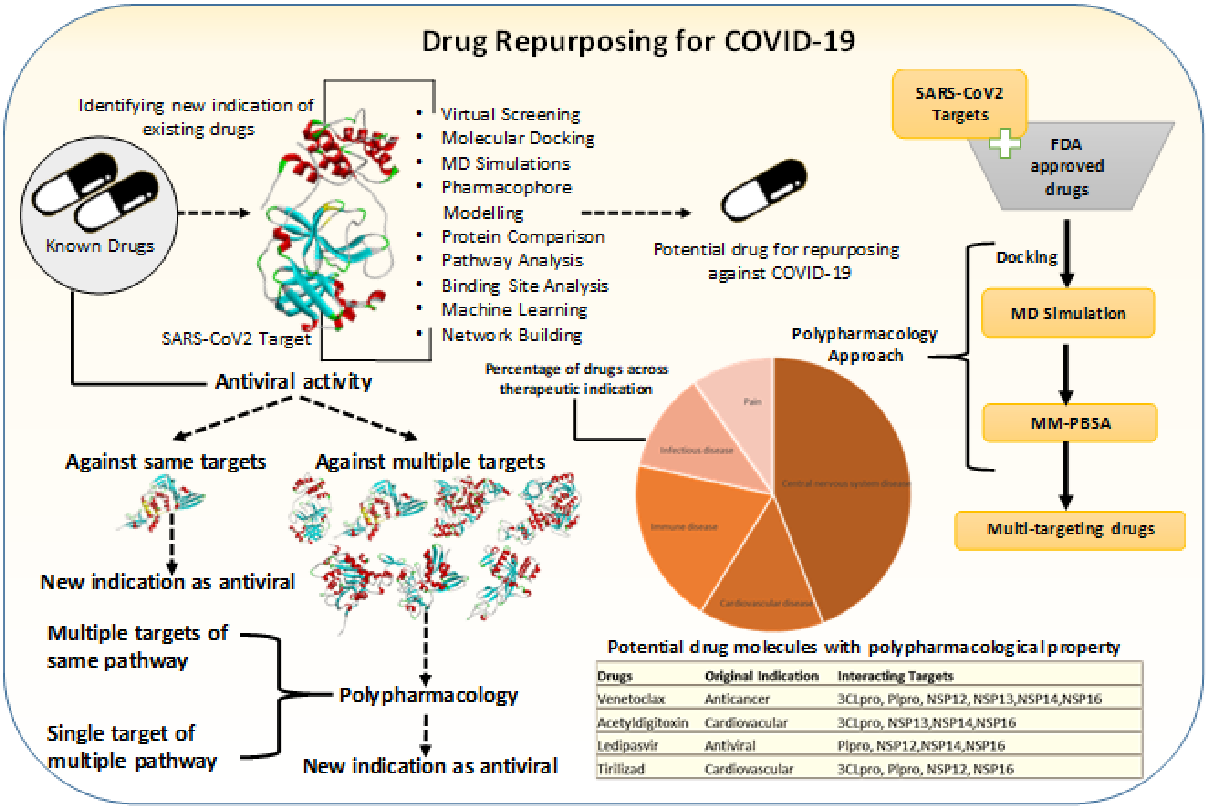


**Figure S3.** An overview of drug repurposing approach for COVID-19 disease which involves identifying of existing drugs with potential therapeutic indications against multiple targets. This may include assessing potential novel uses for drugs based on their known mechanisms of action and interactions with druggable targets, as well as screening existing drugs against the targets in the same pathway or targets from multiple pathways. Applying a unique combination of drug repurposing and polypharmacology approach has led to identification of potential leads that can be repurposed for COVID-19 disease.

**Table S1:** The list of online databases and sources used for the collection of disease specific information for the SARS-CoV-2 disease library in $MPDS^{COVID-19}$.

| Sl. No | Database/Server | Link (Accessed on $12^{th}$ July) |
|---|---|---|
| 1 | PDB | https://www.rcsb.org/ |
| 2 | UniProt | https://www.uniprot.org/ |
| 3 | NCBI | https://www.ncbi.nlm.nih.gov/ |
| 4 | PubMed | https://pubmed.ncbi.nlm.nih.gov/ |
| 5 | BindingDB | https://www.bindingdb.org/rwd/bind/index.jsp |
| 5 | KEGG | http://www.kegg.jp/ or http://www.genome.jp/kegg/ |
| 6 | ChEMBL | https://www.ebi.ac.uk/chembl/ |
| 7 | DrugBank | https://go.drugbank.com/ |
| | **Online Source** | **Link (Accessed on $12^{th}$ July)** |
| 7 | The New York Times | https://www.nytimes.com/interactive/2020/science/coronavirus-drugs-treatments.html |
| | NIH COVID-19 Treatment Guidelines | https://www.covid19treatmentguidelines.nih.gov/ |
| 8 | BMJ | https://www.bmj.com/content/370/bmj.m3379 |
| 9 | GoodRx | https://www.goodrx.com/conditions/covid-19/coronavirus-treatments-on-the-way |
| 10 | World Health Organization (WHO) | https://www.who.int/activities/tracking-SARS-CoV-2-variants |
| 11 | European Centre for Disease Control and Prevention (ECDC) | https://www.ecdc.europa.eu/en/covid-19/variants-concern |
| 12 | FDA Drugs Emergency Preparedness | https://www.fda.gov/drugs/emergency-preparedness-drugs/coronavirus-covid-19-drugs |
| 13 | FDA Press Announcements | https://www.fda.gov/news-events/press-announcements/coronavirus-covid-19-update-fda-authorizes-new-monoclonal-antibody-treatment-covid-19-retains |

**Table S2.** List of AI-Powered drug discovery tools along with its URL, brief description and reference.

| Sl. No | AI-Powered Drug Discovery Tools | URL (Accessed on 10th July 2023) | Description | Reference |
|---|---|---|---|---|
| 1 | AlphaFold | https://alphafold.ebi.ac.uk/ | A DeepMind AI system that predicts 3D structure of protein using the amino acid sequence of protein | 1 |
| 2 | DeepChem | https://deepchem.io/ | A python-based DeepChem provides a set of features for using deep learning to solve challenges in drug development and cheminformatics. | 2 |
| 3 | ODDT (Open Drug Discovery Toolkit) | https://github.com/oddt/oddt | An open-source tool for computer-aided drug discovery. CADD pipelines are created by ODDT using machine learning scoring methods | 3 |
| 4 | Cyclica | https://cyclicarx.com/ | A platform that allows for prioritization of compounds based on their on- and off-target polypharmacology profiles and their ADMET properties | 4 |
| 5 | Exscientia | https://www.exscientia.ai/ | An AI-driven pharmaceutical technology firm dedicated to finding, creating, and developing pharmaceuticals in the quickest and most efficient way. | - |
| 6 | ATOM Modeling PipeLine (AMPL) | https://github.com/ATOMconsortium/AMPL | An open-source, modular, extendable software pipeline enabling model development and sharing for advancing drug discovery | 5 |

**Table S3.** List of available severs and web portals deployed using Galaxy platform.

| Selected servers | URL (Accessed on 10th July 2023) | References |
|---|---|---|
| **Bioinformatics, Drug Discovery and Computational Chemistry** | | |
| Galaxy India | https://galaxyproject.org/use/galaxy-india/ | 6 |
| BIPAA | https://bipaa-galaxy.genouest.org/root/login?redirect=%2F | - |
| MiRGalaxy | https://hub.docker.com/r/glogobyte/mirgalaxy | 7 |
| BRIDGE | https://galaxyproject.org/use/bridge/ | 8 |
| COVID-19 | https://covid19.galaxyproject.org/ | |
| HyPhy HIV NGS Tools | https://galaxyproject.org/use/hyphy/ | 9 |
| FROG | http://frogs.toulouse.inra.fr/ | 10 |
| Anastasia | http://motherbox.chemeng.ntua.gr/anastasia_dev/ | 11 |
| BioBix | http://galaxy.ugent.be/ | 12 |
| BioDivine toolset | https://biodivine-vm.fi.muni.cz/galaxy/ | 13 |
| Graphclust | https://graphclust.usegalaxy.eu/ | 14 |
| MPDS | http://mpds.neist.res.in/ | 15 |
| Chemical ToolBox | https://cheminformatics.usegalaxy.eu/ | 16 |
| GCAC | http://ccbb.jnu.ac.in/gcac | 17 |
| ChemFlow | https://vm-chemflow-francegrille.eu/ | 18 |
| Halogen Bonding | http://134.2.17.68:8081/ | 19 |
| OSDDlinux LiveGalaxy | https://galaxyproject.org/use/osddlinux-livegalaxy/ | 20 |
| Proteogenomics Gateway | https://galaxyproject.org/use/proteogenomics-gateway/ | 21 |
| ProteoRE | https://proteore.org/ | 22 |
| Protologger | http://www.protologger.de/tours/core.galaxy_ui | 23 |
| **Advanced Data Science Tools** | | |
| Machine Learning Workbench | https://galaxyproject.org/use/ml-workbench/ | 24 |
| Galactic Circos | https://github.com/galaxyproject/tools-iuc/tree/master/tools/circos | 25 |
| Interactive environments (RStudio, Jupyter) | https://training.galaxyproject.org/training-material/topics/galaxy-interface/tutorials/rstudio/tutorial.html https://training.galaxyproject.org/training-material/topics/galaxy-interface/tutorials/galaxy-intro-jupyter/tutorial.html | 26 |
| Image analysis | https://training.galaxyproject.org/trainingmaterial/topics/imaging/tutorials/imaging-introduction/tutorial.html | 27 |
| IWC | https://dockstore.org/organizations/iwc | - |
| MCMICRO | https://spatialomics.usegalaxy.eu/ | 28 |
| immuneML | https://galaxy.immuneml.uiocloud.no/ | 29 |
| **Galaxy commercial clouds** | | |
| Globus Genomics | https://galaxyproject.org/use/globus-genomics/ | 30 |
| AnVIL | https://galaxyproject.org/use/anvil/ | 31 |
| PhenoMeNal | https://github.com/phnmnl/container-galaxy-k8s-runtime | 32 |
| Terra | https://galaxyproject.org/use/terra/ | 33 |

**Table S4.** List of publicly available various COVID-19 disease related databases and their description.

| S. No. | Databases | URL (Accessed on 10th July 2023) | Description | Ref |
|---|---|---|---|---|
| 1. | COVID-evidence | https://covid-evidence.org/database | A database for a planned, ongoing and completed trials to treat and prevent COVID-19 | 34 |
| 2. | OxCOVID19 | https://covid19.oii.ox.ac.uk/database/ | A large, single-centre, multimodal relational database. | 35 |
| 3. | LitCOVID | https://www.ncbi.nlm.nih.gov/research/coronavirus/#data-download | A literature hub for tracking up-to-date scientific information. | 36 |
| 4. | ClinicalTrials.gov | https://clinicaltrials.gov/ct2/results?cond=COVID-19 | A database of clinical studies. | 37 |
| 5. | DockCoV2 | https://dockcov2.org/drugs/ | Experimental information about MERS and SARS-CoV. | 38 |
| 6. | Lexicomp Online Drug Database | https://kutuphane.istinye.edu.tr/en/announcements/lexicomp-online-drug-database-covid-19-contents | Drug and clinical information about COVID-19. | 39 |
| 7 | COVID-19 Radiography Database | https://www.kaggle.com/datasets/tawsifurrahman/covid19-radiography-database | A database of COVID-19 patients Chest X-ray images and Lung masks. | 40 |
| 8 | DrugBank | https://go.drugbank.com/covid-19 | A comprehensive, online database containing information on drugs and drug targets. | 41 |
| 9. | SARSCOVIDB | https://sarscovidb.org/ | A database that contains viral strains, hosts, methodologies, genes/proteins with the altered expression. | 42 |
| 10 | GISAID | https://gisaid.org/ | An open access genomic data from all influenza viruses and the coronavirus causing COVID-19. | 43 |
| 11 | ImmuneCODE™ | https://immunerace.adaptivebiotech.com/data/ | A database for decoding the adaptive immune response to COVID19 along with detailed population view. | 44 |
| 12 | CORDITE | https://cordite.mathematik.uni-marburg.de/#/ | A drug interaction database to address viral proteins or human proteins to treat COVID19. | 45 |
| 13 | Coronavirus Antiviral & Resistance Database | https://covdb.stanford.edu/ | A database of antiviral & resistance information. | 46 |
| 14 | DBatVir | http://www.mgc.ac.cn/DBatVir/ | A database of 3873 sequences from coronaviridae family. | 47 |
| 15. | SARS-CoV-2 Database | https://covid19.sfb.uit.no/ | A knowledge database of SARS-CoV-2 virus research compiled from publicly available resources. | 48 |

| | | | | |
|---|---|---|---|---|
| 16. | Therapeutic Target Database | http://db.idrblab.net/ttd/ | A collection of anticoronavirusdrugs and their associated targets and pathways. | 49 |
| 17. | NCBI Virus | https://www.ncbi.nlm.nih.gov/labs/virus/vssi/#/ | A community portal for viral sequence data from RefSeq, GenBank and other NCBI repositories. | 50 |
| 18. | GESS | https://biokeanos.com/source/GESS-db | A database of global evaluation of SARS-CoV-2 sequences. | 51 |
| 19 | IDbSV | http://idbsv.medbiotech-lab.ma/ | A largest repository of SARS-CoV-2 variants and the largest analysis of SARS-CoV-2 genomes. | 52 |
| 20 | COVID-19 Data Portal | https://www.covid19dataportal.org/ | The European COVID-19 Data Platform for data sharing and analysis to accelerate coronavirus research. | 53 |
| 21 | NCATS Open data portal | https://opendata.ncats.nih.gov/covid19/index.html | A collection of datasets by screening a panel of SARS-CoV-2-related assays against all approved drugs. | 54 |
| 22 | The COVID-19 Drug and Gene Set Library | https://maayanlab.cloud/covid19/ | A collection of drug and gene sets related to COVID-19 research. | 55 |
| 23 | National Institute of Allergy and Infectious Diseases (NIAID) | https://www.niaid.nih.gov/research/accessing-clinical-data | A cloud-based, secure data platform that enables sharing of and access to reports and data sets related to COVID-19. | 56 |
| 24 | Nextstrain | https://nextstrain.org/ncov/gisaid/global/6m | An open-source database to empower the wider genomic epidemiology and public health. | 57 |
| 25 | SARS-CoV-2 related structures | https://covid19.bioreproducibility.org/ | A database with 2465 SARS-CoV-2 protein structures and 143 additional structures of other corona viruses. | 58 |
| 26 | ERACODA | https://www.eracoda.org/ | A European database collecting clinical information of patients on kidney replacement therapy with COVID-19. | 59 |
| 27 | Virus-CKB | https://www.cbligand.org/g/virus-ckb | A knowledgebase for COVID-19 and similar viral infection research. | 60 |
| 28 | CoronaVR | https://bioinfo.imtech.res.in/manojk/coronavr/ | A database that contains potential epitopes, therapeutics, etc. | 61 |
| 29 | Drug Gene Interaction database | https://www.dgidb.org/ | A database of drug-gene interactions and the druggable genome, mined from over thirty trusted sources. | 62 |
| 30 | STRING Database | https://string-db.org/ | A database of known and predicted protein-protein interactions. | 63 |

| | | | | |
|---|---|---|---|---|
| 31 | COVID-19 Genome Sequence Dataset | https://registry.opendata.aws/ncbi-covid-19/ | A centralized sequence repository for all strains of novel corona virus (SARS-CoV-2). | 64 |
| 32 | COVID-19 Molecular Structure and Therapeutics Hub | https://covid.molssi.org/ | A community data repository for structure, models, therapeutics, simulations related computations for COVID-19. | 65 |
| 33 | COVID19db | http://www.biomedical-web.com/covid19db/home | A comprehensive platform to discover potential drugs and targets of COVID-19 at whole transcriptome scale. | 66 |
| 34 | H2V | http://www.zhounan.org/h2v/ | A knowledge base of human genes/proteins that respond to the infection of SARS-CoV-2, SARS-CoV and MERS-CoV. | 67 |
| 35 | SARS-CoV-2 Virion and Proteins | https://3dprint.nih.gov/niaid/sars-cov-2 | A database consists of structures released by the Protein Data Bank, as well as additional “Featured” structures curated by NIAID, in 3D-printable formats. | 68 |
| 36 | MIDAS | https://midasnetwork.us/covid-19/ | A collection of more than 300 digital resources relevant to COVID modeling. | 69 |
| 37 | outbreak.info | https://outbreak.info/situation-reports#voc | A database with 12,742,821 sequences from GISAID. | 70 |
| 38 | COVIDium | http://kraza.in/covidium/ | A list of all the relevant databases characterized into 10 broad categories. | 71 |
| 39 | SCoV2-MD | https://submission.gpcrmd.org/covid19/home/ | Share, visualize and analyze MD data of SARS-CoV-2 related proteins | 72 |
| 40 | Comparative Toxicogenomics Database | http://ctdbase.org/detail.go?type=disease&acc=MESH%3AD000086382 | A database of manually curated information about chemical–gene/protein interactions, chemical–disease and gene–disease relationships. | 73 |
| 41 | REACTOME | https://reactome.org | An open-source, open access, manually curated and peer-reviewed pathway database. | 74 |
| 42 | ViPR | https://www.viprbrc.org/brc/home.spg?decorator=corona_ncov | A database with latest SARS-CoV-2 Variants and Lineages of Concern. | 75 |
| 43 | INSDC | https://ncbiinsights.ncbi.nlm.nih.gov/2020/08/17/insdc-covid-data-sharing/ | A database that capture, organize, preserve and present nucleotide sequence data. | 76 |
| 44 | COVID-19 Drug Repurposing Database | https://www.excelra.com/covid-19-drug-repurposing-database/ | An open-access database of ‘Approved’ small molecules and biologics, which can rapidly enter either Phase 2 or 3, or may even be used directly in clinical settings against COVID-19. | 77 |
| 45 | NCBI SARS-CoV-2 Resources | https://www.ncbi.nlm.nih.gov/sars-cov-2/ | A comprehensive information from NCBI. | 78 |

| | | | | |
|---|---|---|---|---|
| 46 | CAS COVID-19 antiviral candidate compounds dataset | https://www.cas.org/covid-19-antiviral-compounds-dataset | An open-source dataset with 50,000 chemical substances includes antiviral drugs and related compounds. | 79 |
| 47 | BioExcel-CV19 | https://bioexcel-cv19.bsc.es/#/ | A web-access to atomistic-MD trajectories for macromolecules involved in the COVID-19. | 80 |
| 48 | Rfam | https://rfam.org/covid-19 | A database of new and updated Coronavirus families | 81 |
| 49 | Malacard | https://www.malacards.org/card/covid_19 | An integrated GeneCards database of human genes. | 82 |
| 50 | National Genomics Data Center | https://ngdc.cncb.ac.cn/ | An open access database with big genomics data. | 83 |
| 51 | CoV-AbDab | http://opig.stats.ox.ac.uk/webapps/covabdab/ | A database of immune response to SARS-CoV-2 infection and vaccination. | 84 |
| 52 | COVIDep | https://covidep.ust.hk/ | A database for B-cell and T-cell epitopes that can serve as potential vaccine targets for COVID-19. | 85 |
| 53 | VirHostNet | https://virhostnet.prabi.fr/ | It is a bioinformatics information system dedicated to the bio-curation, data integration, reproducible systems-level analysis and visualization of viral protein-protein networks. | 86 |
| 54 | GENCODE | https://www.gencodegenes.org/human/covid19.html | It contains annotation of human protein-coding genes linked to SARS-CoV-2 infection. | 87 |
| 55 | RCoV19 | https://ngdc.cncb.ac.cn/ncov/ | A database of comprehensive integration of genomic and proteomic sequences as well as their metadata information from the GISAID, NCBI, NMDC and CNCB/NGDC | 88 |
| 56 | COVID-19 UniProtKB | https://covid-19.uniprot.org/uniprotkb?query=* | It has latest available pre-release UniProtKB data for the SARS-CoV-2 coronavirus and other entries relating to the COVID-19 outbreak. | 89 |
| 57 | COG-UK-Mutation Explorer | https://sars2.cvr.gla.ac.uk/cog-uk/ | It provides information and structural context on mutations and associated variants in the genes encoding SARS-COV-2 proteins | 90 |
| 58 | ViralZone | https://viralzone.expasy.org/ | It provides resources related to beta-coronavirus genome, proteome, interactome, coronavirus life cycle, antiviral drugs and treatment. | 91 |
| 59 | CoV-GLUE | https://cov-glue.cvr.gla.ac.uk/ | It has the information of mutations, insertions and deletions which have been | 92 |

| | | | | |
|---|---|---|---|---|
| | | | observed in GISAID hCoV-19 sequences. | |
| 60 | Cellosaurus | https://web.expasy.org/cellosaurus/sars-cov-2.html | A knowledge resource on cell lines for SARS-CoV-2. | 93 |
| 61 | Corona OMA Browser | https://corona.omabrowser.org/oma/home/ | A database that encompasses the human SARS viruses, and other viruses. | 94 |
| 62 | GlyConnect | https://glyconnect.expasy.org/covid-19 | Provide information on characterizing the molecular components of protein glycosylation for SARS-CoV-2. | 95 |
| 63 | neXtProtD | https://www.nextprot.org/ | It includes information about human proteins that bind SARS-CoV-2. | 96 |
| 64 | SWISS-MODEL | https://swissmodel.expasy.org/repository/species/2697049 | Provides 3D homology models and links to experimental structures in PDB for SARS-CoV-2 proteins. | 97 |
| 65 | Bioinformatics Resources for SARS-CoV-2 | https://www.clinbioinfosspa.es/CovidResources | It provides links to bioinformatics resources useful to track the evolution and progression of the SARS-CoV-2. | 98 |
| 66 | CoV2ID | http://covid.portugene.com/cgi-bin/COVid_home.cgi | This database has updated list of oligonucleotides for SARS-CoV-2 | 99 |
| 67 | canSAR | https://corona.cansar.icr.ac.uk/ | A COVID-19 drug discovery platform. | 100 |
| 68 | C-I-TASSER | https://seq2fun.dcmb.med.umich.edu//COVID-19/ | Provides 3D structural models and function annotation for all proteins encoded by the genome of SARS-CoV-2. | 101 |
| 69 | CoVex | https://exbio.wzw.tum.de/covex/ | It integrates virus-human interactions for SARS-CoV-2 and SARS-CoV-1. | 102 |
| 70 | DBCOVP | http://covp.immt.res.in/ | Database of coronavirus virulent glycoproteins | 103 |
| 71 | SARS-CoV-2 Proteome-3D | https://sars3d.com/ | Annotated S SARS-CoV-2-3D proteome database. | 104 |
| 72 | hCoronavirusesDB | http://hcoronaviruses.net/#/ | An integrated bioinformatics resource for human coronaviruses. | 105 |
| 73 | COVIDOUTCOME | https://www.covidoutcome.com/ | A database that informs about the severity based on Mutations in the SARS-CoV-2 Genome. | 106 |
| 74 | Chemical Checker | https://sbnb.irbbarcelona.org/covid19/ | A database of small molecules with similar chemical and bioactivity features to the reported drugs in a universe of 800 thousand bioactive compounds | 107 |
| 75 | Signor | https://signor.uniroma2.it/covid/index.php?beta=3.0 | A database in which every entry point has experimental evidence. With available evidence, it has | 108 |

| | | | | |
|---|---|---|---|---|
| | | | relevant information for the COVID-19 pathology. | |
| 76 | Human Protein Atlas | https://www.proteinatlas.org/humanproteome/sars-cov-2 | A database for knowledge-base of disease and efforts to develop diagnostic tools and therapeutic drugs to combat the pandemic. | 109 |
